# Artemin sensitises mouse (*Mus musculus*) and naked mole-rat (*Heterocephalus glaber*) sensory neurons *in vitro*

**DOI:** 10.1101/2025.04.10.648171

**Authors:** Lanhui Qiu, Ewan St. John Smith

**Author notes:** To whom correspondence should be addressed: Prof. Ewan St. John Smith. Authorship statement LQ designed studies, conducted experiments, analyzed data and wrote the initial draft. EStJS supervised the project, edited the manuscript and acquired funding.

## Abstract

The naked mole-rat (NMR, *Heterocephalus glaber*) is a subterranean rodent that exhibits a range of unusual physiological traits, including diminished inflammatory pain. For example, nerve growth factor (NGF), a key inflammatory mediator, fails to induce sensitization of sensory neurons and thermal hyperalgesia in NMRs. This lack of NGF-induced neuronal sensitization and thermal hyperalgesia results from hypofunctional signaling of the NGF receptor, tropomyosin receptor kinase A (TrkA). Like NGF-TrkA signaling, the neurotrophic factor artemin, a member of the glial cell line-derived neurotrophic factor (GDNF) family, is implicated in mediating inflammatory pain through its receptor, GDNF family receptor α3 (GFRα3), which is expressed by a subset of dorsal root ganglia (DRG) sensory neurons. Here we investigated GFRα3 expression in DRG neurons of mice and NMRs, as well as measuring the impact of artemin on DRG sensory neuron function in both species *in vitro*. Using immunohistochemistry, we observed a similar abundance of GFRα3 in mouse and NMR DRG sensory neurons, high coexpression with the transient receptor potential vanilloid 1 (TRPV1) ion channel suggesting that these neurons are nociceptive neurons. Using *in vitro* electrophysiology to record from cultured DRG sensory neurons, we observed that artemin induced depolarization of the resting membrane potential and decreased the rheobase in both species, as well as diminishing the degree of TRPV1 desensitization to multiple capsaicin stimuli. Overall, results indicate that artemin similarly sensitizes sensory neurons in both mice and NMRs, future *in vivo* studies being required to confirm if the conserved *in vitro* sensitization also occurs *in vivo*.

## 1. Introduction

Nociception functions as a detection and protection mechanism and occurs in both invertebrates and vertebrates, thus demonstrating its fundamental importance in the strive for survival (Smith and Lewin 2009; Sneddon 2018). Whereas many species display nocifensive behaviors to noxious mechanical, thermal, and chemical stimuli, certain species show adaptations that enable them to cause or avoid nociception and provide a survival benefit. For example, the scarlet velvet ant (*Dasymutilla occidentalis*) produces a peptide that is innocuous in mice, but activates invertebrate nociceptive neurons, those sensory neurons that transduce noxious stimuli, and enables an effective defense against attacks from the predatory Chinese mantis *Tenodera sinensis* (Borjon et al. 2025). By contrast, overexpression of the leak channel NALCN (sodium leak channel, nonselective) in sensory neurons of the highveld mole-rat (*Cryptomys hottentotus pretoriae*) prevents nociceptive neuron activation by the sting of the Natal droptail ant (*Myrmicaria natalensis*) (Eigenbrod et al.). The naked mole-rat (NMR, *Heterocephalus glaber*) shows behavioral insensitivity to acid, capsaicin, and diminished inflammatory pain (Park et al. 2008), as well as a plethora of adaptations that support healthy ageing to 35+ years of age (Buffenstein et al. 2022). The behavioral acid insensitivity of NMRs appears to arise from genetic variation in the voltage-gated sodium channel Na_V_1.7 (Smith et al. 2011), whereas capsaicin insensitivity occurs due to altered spinal cord sensory neuron wiring, rather than an alteration in primary afferent neuron activity (Park et al. 2008). In addition, the inability of the inflammatory mediator nerve growth factor (NGF) to induce thermal hyperalgesia in NMRs is due to hypofunctionality of the NGF receptor, tropomyosin receptor kinase A (TrkA) (Omerbašić et al. 2016).

However, NGF is not the only neurotrophic factor implicated in inflammatory pain. Artemin is a member of the glial cell line-derived neurotrophic factor (GDNF) family, which signals through its receptor, GDNF family receptor α3 (GFRα3), and contributes to nociceptive neuron sensitization and hyperalgesia (Elitt et al. 2006; Malin et al. 2006; Morgan et al. 2023). Artemin’s receptor, GFRα3, is expressed by a significant proportion of nociceptive neurons (Zeisel et al. 2018; Hockley et al. 2019), including those innervating the knee joint (Morgan et al. 2023), making artemin and GFRα3 potential therapeutic targets for joint pain. Indeed, in animal models of inflammatory and osteoarthritis pain, blocking artemin signaling with neutralizing antibodies reduces nociceptive neuron sensitization and alleviates pain (Nencini et al. 2018, 2019; Minnema et al. 2022; Morgan et al. 2023). Similarly, anti-GFRα3 antibody treatment has proved efficacious in mouse joint pain models, but no benefit was observed in a Phase 2 study of osteoarthritis pain in humans (Somersan-Karakaya et al. 2023). Findings from transgenic mouse models indicate that artemin directly sensitizes nociceptive neurons by influencing activity of the transient receptor potential cation channel subfamily V member 1 (TRPV1) ion channel, such that mice overexpressing artemin show heightened responses to the TRPV1 agonist capsaicin (Elitt et al. 2006). Similarly, Malin et al. (2006) reported that artemin enhances DRG neuron sensitivity to capsaicin, showing its role in modulating TRPV1 activity and nociceptive neuron excitability. Considering artemin’s ability to sensitize nociceptive neurons and the involvement of artemin-GFRα3 signaling in inflammatory pain, we set out to determine how artemin modulates neuronal function in NMRs, a species that exhibits decreased inflammatory pain.

Conducting whole-cell patch-clamp electrophysiology, we demonstrate that artemin sensitizes both mouse and NMR sensory neurons with GFRα3 expression being similar across species, results which would suggest that artemin-mediated pathways are conserved between mice and NMRs. These findings offer new insights into the conservation of nociceptive mechanisms across species at the *in vitro* level, future *in vivo* studies being required to determine if artemin causes behavioral hypersensitivity in NMRs as has been previously demonstrated in mice.

## 2. Materials and methods

### 2.1 Animals

All animal experiments were performed in accordance with the Animals (Scientific Procedures) Act 1986 Amendment Regulations 2012 under Project Licenses granted to E.St.J.S. (P7EBFC1B1 and PP5814995) by the Home Office and approved by the University of Cambridge Animal Welfare Ethical Review Body. Experiments were performed using a mixture of male and female C57BL6/J mice (10-15 weeks old) and a mixture of male and female, non-breeder/subordinate, NMRs (23-145 weeks old). Age matching mice and NMRs is not straightforward, but considering the long-lifespan of NMRs vs. mice, all animals used in this study can be considered young adults. Mice were purchased from Envigo and housed conventionally with nesting material and a red plastic shelter in a temperature-controlled room at 21 °C, with a 12-hour light/dark cycle and access to food and water ad libitum. NMRs were bred in-house and maintained in an interconnected network of cages in a humidified (∼55%) temperature-controlled room at 28-32 °C with red lighting (08:00−16:00). NMRs had access to food ad libitum. NMRs used in this study came from five different colonies. Animals were humanely killed by CO_2_ exposure followed by cervical dislocation (mice) or decapitation (NMRs). To limit the total number of animals used, DRG were taken from each animal for both immunohistochemistry (IHC) and electrophysiological analysis.

### 2.2 DRG neuron isolation and culture

DRG neuron isolation and culture were performed as previously described (Ai et al. 2023; Pattison et al. 2024). In brief, before collecting DRG, poly-D-lysine glass-bottomed dishes (MatTek, USA) were coated with laminin (1mg/ml, Invitrogen UK, 23017-015) and incubated at 37°C for 1 hour before removing excess laminin, washing with H_2_O and drying. Lumbar DRG (L2-L5, i.e. those that predominantly innervate the knee joint (da Silva Serra et al. 2016; Chakrabarti et al. 2020) were collected post-mortem and placed into cold dissociation medium (L-15 Medium (Gibco™ UK, 31415-086) with 1.5% NaHCO_3_ (Gibco™ UK, 25080-094)). DRG were enzymatically digested in prewarmed collagenase solution (dissociation medium with 1 mg/ml collagenase (Sigma-Aldrich UK, C9891) and 6 mg/ml bovine serum albumin (BSA, Sigma-Aldrich UK, A2153)) for 15 mins followed by digestion in trypsin solution (dissociation medium with 1 mg/ml trypsin (Sigma-Aldrich UK, T9935), 6 mg/ml BSA, Sigma, UK) for 30 mins at 37°C before mechanical trituration with a P1000 tip 20-25 times in culture medium (dissociation medium with 1% penicillin-streptomycin (Gibco UK, 15140-122) + 10% fetal bovine serum (Sigma-Aldrich UK, F7524). Following trituration, brief centrifugation (1000 g, 30s) was used to separate remaining DRG tissue from dissociated neurons. The resulting supernatant, containing the dissociated neurons, was collected in a separate tube, and then 2 ml of culture medium was added to pelleted DRG tissues for further trituration. Trituration and brief centrifugation were repeated 5 times until 10 ml of supernatant was collected. Collected supernatant was centrifuged at 1000 g for 5 mins and the pellet was resuspended in culture medium and 100 µl plated into the center of each glass-bottomed dish. Neurons were incubated at 37°C with 5% CO_2_for 2 hours before then adding 1.9 mL of culture medium; neurons were maintained in the incubator for 18–24 hours before electrophysiological recordings.

### 2.4 Immunohistochemistry

Immunohistochemistry was performed as previously described (Chakrabarti et al. 2020). Briefly, DRG were collected post-mortem and post-fixed for 1h in 4% paraformaldehyde (PFA), followed by overnight incubation in 30% (w/v) sucrose at 4 °C for cryoprotection. DRG were next embedded in Shandon M-1 Embedding Matrix (Thermo Fisher Scientific, 10056778), snap frozen in 2-methylbutane (Honeywell International, M32631) on dry ice and stored at -80 °C. Embedded DRG were sectioned (12μm) using a Leica Cryostat (CM3000; Nussloch, Germany) on to Superfrost Plus microscope slides (Thermo Fisher Scientific, MIC30220) and stored at -20°C until staining.

Slides were defrosted, washed with phosphate-buffered saline (PBS)-tween and blocked in antibody diluent solution: 0.2% (v/v) Triton X-100, 5% (v/v) donkey serum and 1% (v/v) bovine serum albumin in PBS for 1-h at room temperature before overnight incubation at 4°C with anti-TRPV1 (1:500, guinea pig polyclonal, Alomone Labs, ACC-030-GP) and anti-GFRα3 (1:300, goat polyclonal R&D systems, AF2645) primary antibodies in antibody diluent. Slides were washed three times with PBS-Tween and incubated with anti-guinea pig AF594 (Invitrogen, A11076) and anti-goat Alexa-568 (Invitrogen, A11057) secondary antibodies for 2 hours at room temperature. Slides were washed in PBS-Tween, mounted and imaged with an Olympus BX51 microscope (Tokyo, Japan) and QImaging camera (Surrey, Canada). Exposure levels were kept constant for each slide, and contrast enhancements were made the same for all slides. Negative controls without the primary antibody showed no staining.

For analysis, sections were blinded to experimental groups, and ImageJ software (version 1.53k) was used. An automatic “minimum error” threshold algorithm was applied to 8-bit images to distinguish objects from the background. Neurons within each DRG section were manually selected, and their mean grey values were measured and normalized to the range defined by the lowest and highest intensity neurons in that section. A neuron was classified as positive for the stain if its normalized grey value exceeded the global minimum across all sections plus 2 standard deviations (SD).

### 2.5 Electrophysiology

Recordings from DRG neurons took place after 15 min incubation with 0.5 μg/ml isolectin B4 (IB4)-Alexa 488 (Invitrogen I21411) or IB4-Alexa Fluor 568 (Invitrogen I21412). Glass patch pipettes (3-6 MΩ, Hilgenberg) were pulled by a P-97 Flaming/Brown puller (Sutter Instruments, USA) from borosilicate glass capillaries and loaded with intracellular solution (ICS), which (in mM) contained KCl (110), NaCl (10), MgCl_2_ (1), EGTA (1), and HEPES (10), adjusted to pH 7.4 with KOH and osmolality adjusted to 300-310 mOsm using sucrose. DRG neurons were bathed in extracellular solution (ECS) (in mM): NaCl (140), KCl (4), CaCl_2_ (2), MgCl_2_ (1), glucose (4), HEPES (10), adjusted to pH 7.4 with NaOH, and osmolality was adjusted to 280-295 mOsm using sucrose. Cells were observed using an Olympus IX70 Fluorescence Microscope. Solutions were applied with a gravity-driven multi-barrel perfusion system (Automate Scientific). Data were acquired with a Multiclamp 700A amplifier (Molecular Devices, USA) and digitized via Digidata 1440A digitizer (Molecular Devices, USA).

For current-clamp recordings, bridge-balance compensation was used to reduce steady-state voltage errors. Data were sampled at 20kHz and filtered at 5kHz. Step current (-100 pA to 2000 pA) for 80 ms through 21 steps or no current was injected to generate action potentials (APs) under current-clamp mode. For voltage-clamp recordings, solutions were applied using a gravity-driven multi-barrel perfusion system. All recordings were made using an Multiclamp 700A amplifier in combination with Clampex (Molecular Devices, USA) software. Pipette and membrane capacitance were compensated, with series resistance being compensated by ∼70% to minimize voltage errors. IB4-positive neurons were visualized by observation of Alexa-488 or Alexa-568 fluorescence (CellCam Kikker 100MT)

*Escherichia coli* derived mouse artemin (Ala112-Gly224, R&D Systems, 1085-AR) was used to assess artemin’s ability to sensitize DRG neurons. Baseline intrinsic excitability was first measured using current-clamp recordings, in which action potential parameters, including resting membrane potential (RMP), rheobase, peak amplitude, afterhyperpolarization (AHP) duration, and AHP amplitude, were analyzed with Clampex (Molecular Devices, USA). In the same recordings, TRPV1 function was assessed in voltage clamp configuration by applying 1 μM capsaicin (Sigma-Aldrich, M2028) for 5 seconds. To evaluate the sensitizing effects of artemin, neurons were exposed to a 7-minute perfusion of 100 ng/ml artemin, before repeating current-clamp, voltage-clamp and capsaicin sensitivity recordings; in control experiments, neurons were exposed to ECS for 7-minutes.

### 2.6 Statistics

Data were assessed for normality using the Shapiro-Wilk test, and parametric or non-parametric statistics were used as appropriate. IHC results are reported as mean ± standard error of the mean (SEM) Patch clamp data were analyzed using paired or unpaired t-tests as appropriate. Results are reported as mean ± SEM. Statistical analysis and graph generation was carried out in GraphPad Prism 8.0 software (USA), Biorender®, R studio® and Python.

## 3. Results

### 3.1 High coexpression of TRPV1 and GFRα3 in mouse and NMR DRG neurons

To probe the coexpression of GFRα3 with TRPV1, a marker of putative nociceptive neurons, double-labeling immunohistochemistry was performed using lumbar DRG (Fig. 1). In NMR (N=5), 27.80±1.94% of DRG neurons were found to express GFRα3, 27.60±2.29% expressed TRPV1, and 23.60±2.27% expressed both, 84.89% of TRPV1 +ve neurons, i.e. putative nociceptive neurons, expressing GFRα3 (Fig. 1a). In mice (N=5) similar observations were made, 30.57±1.41% of DRG neurons expressing GFRα3, 26.00±1.92% expressing TRPV1, 23.96±1.65% expressing both TRPV1 and GFRα3, with 93.7% of TRPV1 +ve neurons expressing GFRα3 (Fig. 1b). Therefore, in both mice and NMR, most GFRα3-positive neurons coexpress TRPV1. These immunohistochemical data support the hypothesis that in NMR, as in mouse, a subset of putative nociceptive neurons express GFRα3 and thus could subserve sensitizing responses to artemin.

**Fig. 1.**
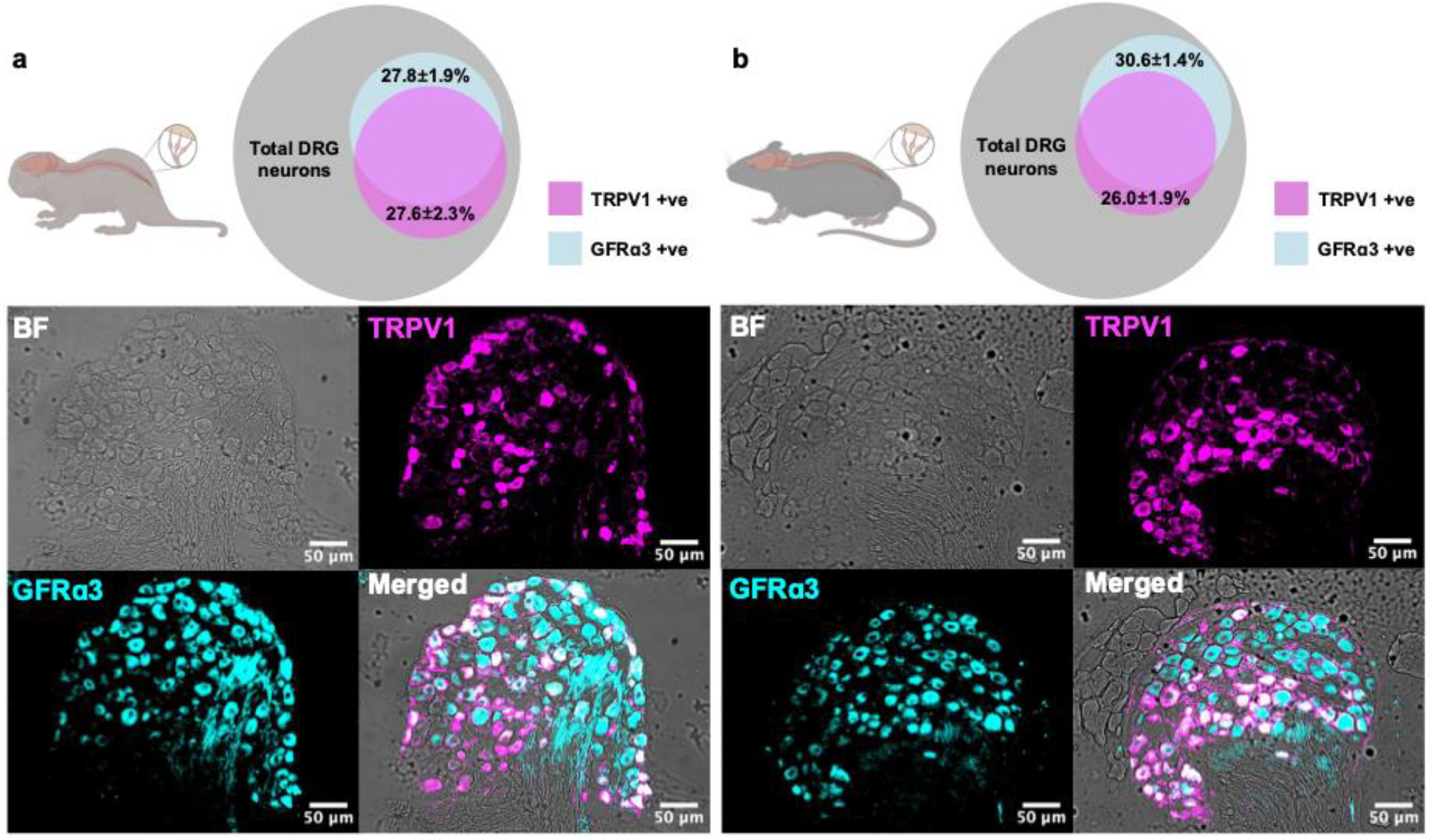
TRPV1 and GFRα3 show high coexpression in both mouse and NMR DRG neurons. (a) In NMR (N=5, n=21 sections, n=1302 cells), 27.80±1.94% of DRG neurons express GFRα3, 27.60±2.29% express TRPV1, and 23.60±2.27% express both. 84.89% of TRPV1 +ve neurons also expressing GFRα3. (b) In mice (N=5, n=23 sections, n=1564 cells), 30.57±1.41% of DRG neurons express GFRα3, 26.00±1.92% express TRPV1, 23.96±1.65% express both, and 93.7% of TRPV1 +ve neurons also expressing GFRα3. Scale bar in all panels, 50 µm. Data are reported as mean±SEM.

### 3.2 Artemin sensitises mouse and NMR DRG neurons

We made whole-cell patch-clamp recordings from isolated mouse and NMR DRG sensory neurons following live-labelling with fluorescently conjugated IB4. IB4 predominantly binds to non-peptidergic small-diameter sensory neurons in mice, while TrkA immunoreactivity is observed more frequently in peptidergic sensory neurons that do not bind IB4 (Averill et al. 1995); IB4 binding and TrkA expression also define largely distinct populations in NMR DRG neurons (Omerbašić et al. 2016). Moreover, single-cell transcriptomic sequencing analyses of mouse DRG neurons show that GFRα3 expression is largely within the TrkA-positive DRG neuron population, and that these neurons often coexpress TRPV1 (Zeisel et al. 2018; Hockley et al. 2019). Therefore, by focusing our analysis on IB4-negative neurons, we aimed to enhance the likelihood that neurons expressed GFRα3 and thus had capacity to respond to artemin, as well as being likely to also express TRPV1; see also Figure 1. Sequence analysis reveals that the NMR GFRα3 receptor shares approximately 74% amino acid identity with mouse GFRα3, indicating a high degree of evolutionary conservation of the receptor between these species; NCBI GFRα3 reference sequences: mouse, NP_034410.3 and NMR, EHB06152.1.

The ability of artemin to sensitize DRG neurons was measured by firstly examining the impact of an acute, 7-minute, incubation of 100 ng/ml artemin on neuronal intrinsic excitability. In DRG neurons isolated from both mice and NMRs, artemin treatment resulted in a reduction in the rheobase and depolarization of the resting membrane potential (Fig. 2 and Table 1). In NMR DRG neurons, artemin caused the rheobase to decrease from 305.33 ± 30.30 pA to 214.22 ± 24.41 pA (Fig. 2a, p<0.01), while the resting membrane potential moved in a depolarizing direction from -48.59 ± 0.92 mV to - 43.84 ± 0.94 mV (Fig. 2b, p<0.0001); in control experiments, 7-minute perfusion of extracellular solution (ECS) did not significantly alter rheobase or resting membrane potential (Fig. 2a-b). Similar effects were observed in mouse DRG neurons, artemin reducing the rheobase from 363.42 ± 40.62 pA to 214.22 ± 41.68 pA (Fig. 2f, p<0.001) and depolarizing the resting membrane potential from - 55.90 ± 0.65 mV to -45.44 ± 1.35 mV (Fig. 2g, p<0.0001), control experiments with ECS perfusion causing no significant effects (Fig. 2f-g).

**Table 1:**
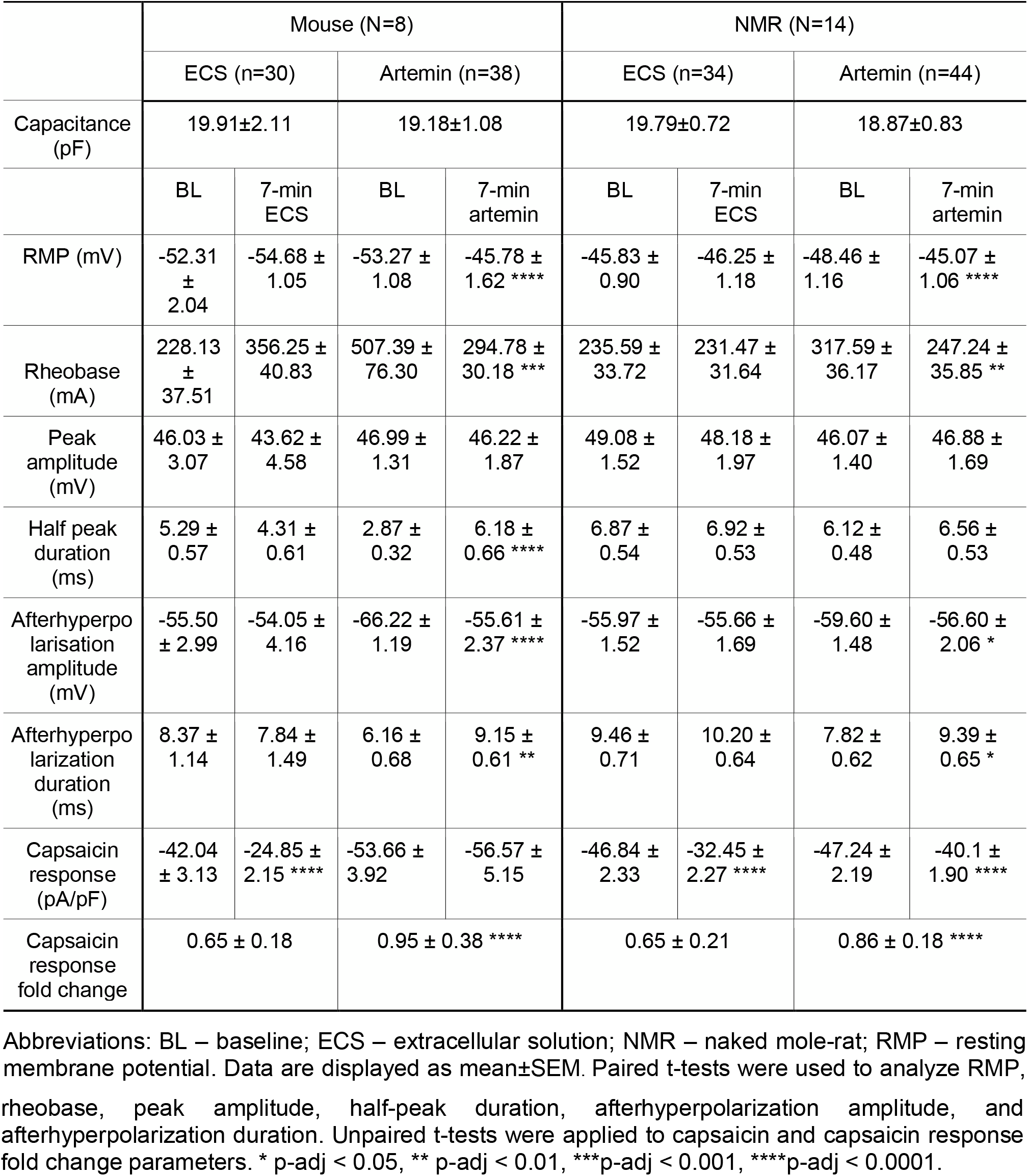
Intrinsic and active properties of DRG neurons.

**Fig. 2.**
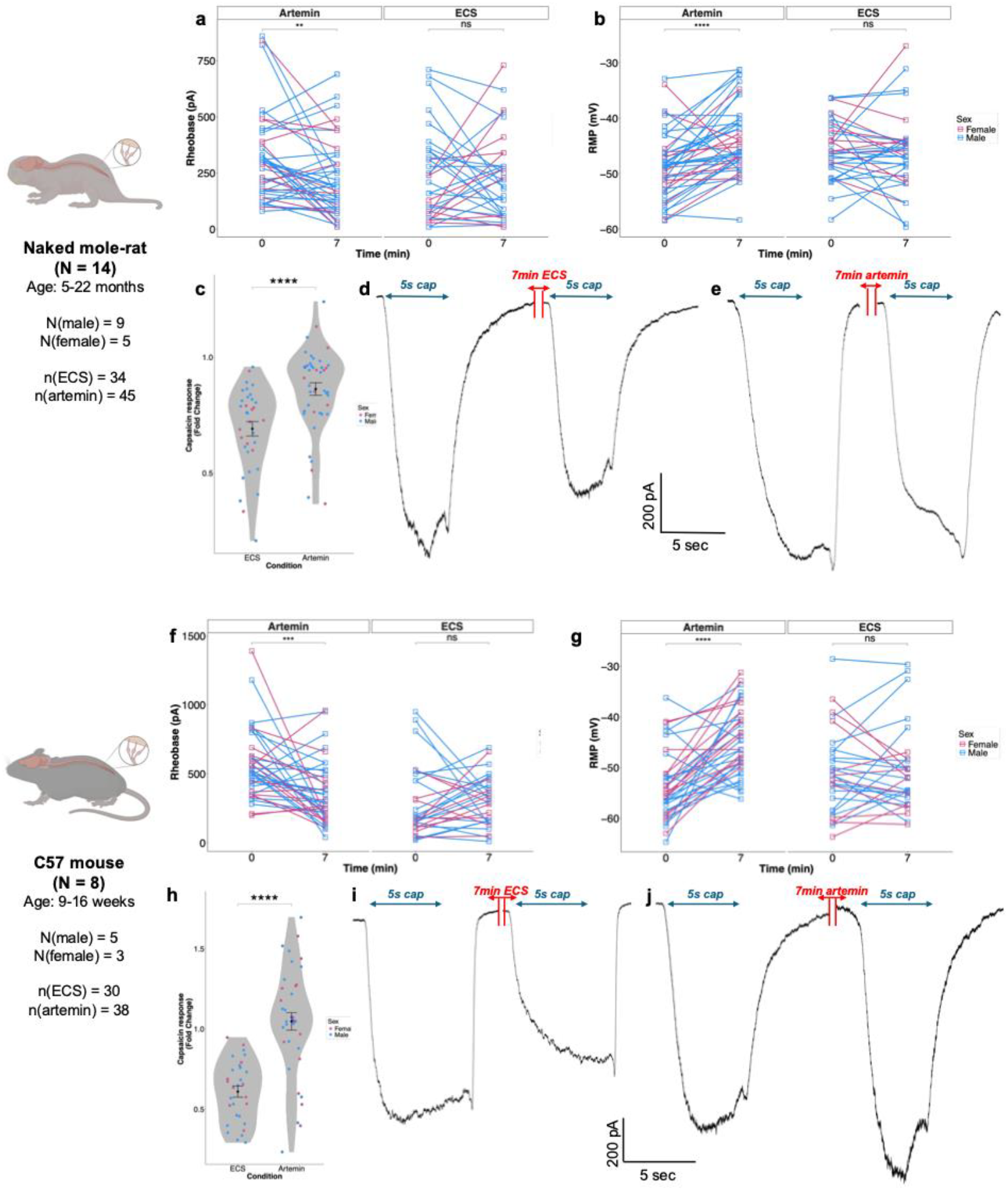
Artemin sensitizes mouse and NMR DRG neurons. (a–c) In NMR DRG neurons, 7-minute perfusion with 100 ng/ml artemin significantly decreased rheobase (a), depolarized the resting membrane potential (b), and diminished TRPV1 desensitization (c-e). Representative traces are shown in (d-e). In mouse DRG neurons, 7-minute perfusion with 100 ng/ml artemin significantly decreased rheobase (f), depolarized the resting membrane potential (g), and diminished TRPV1 desensitization (h-j). Paired t-tests were used to analyze RMP, rheobase, peak amplitude, half-peak duration, afterhyperpolarization amplitude, and afterhyperpolarization duration. Unpaired t-tests were applied to capsaicin response fold change analysis. * p-adj < 0.05, ** p-adj < 0.01, ***p-adj < 0.001, ****p-adj < 0.0001.

In the same recordings, we also examined the ability of artemin to sensitize the function of TRPV1, which has previously been shown to occur in mouse DRG neurons using Ca^2+^-imaging (Malin et al. 2006). Repeated exposure of mouse and NMR DRG neurons to 1μM capsaicin results in desensitization (Omerbašić et al. 2016), which we also observed here. Exposure of mouse or NMR DRG neurons to two capsaicin stimuli 7-minutes apart from each other (Fig. 2d and 2i) resulted in a significant reduction in peak current density from -42.04 ± 3.13 pA/pF to -24.85 ± 2.15 pA/pF in mouse DRG neurons (p<0.0001) and from -46.84 ± 2.33 pA/pF to -32.45 ± 2.27 pA/pF in NMR DRG neurons (p<0.0001). Following exposure to 100 ng/ml artemin for 7-minutes between capsaicin stimulations ameliorated this desensitization in both mouse and NMR DRG neurons (Fig. 2e and 2j), such that in mouse DRG neurons the reduction in peak current density was no longer significant (-53.66 ± 3.92 pA/pF vs. . -56.57 ± 5.15 pA/pF), whereas in NMR DRG neurons the second capsaicin stimulation was still significantly smaller than the first (-46.84 ± 2.33 pA/pF vs. -40.1 ± 1.90 pA/pF, p<0.0001). However, when examining the degree of desensitization in all conditions by calculating the fold change, in NMR neurons, the fold change was 0.69 ± 0.03 for ECS and 0.86 ± 0.02 for artemin (Fig. 2c, p<0.0001). Similarly, in mouse DRG neurons, the fold change was 0.61 ± 0.03 for ECS and 1.05 ± 0.05 for artemin (Fig. 2h, p<0.0001). Thus, although artemin causes a more robust prevention of desenstiization in mouse DRG sensory neurons, artemin has a similar, albeit smaller, effect in NMR DRG sensory neurons.

## 4. Discussion

Evolutionary pressure has resulted in conservation of the mechanisms underpinning nociceptive neuron activity and nocifensive behaviors (Smith and Lewin 2009; Sneddon 2018; Walters et al. 2023). However, some species have evolved adaptations to their environments, which result in unusual nociceptive neuron function and nocifensive behaviors, including NMRs (Smith et al. 2020). Here we made a comparative characterization of the ability of the neurotrophic factor artemin to modulate neuronal activity in mouse and NMR sensory neurons *in vitro*, as well as investigating sensory neuron expression of the receptor artemin, GFRα3.

Artemin is a well-characterized modulator of nociceptive signaling, binding of artemin to GFRα3 activating intracellular signaling cascades that evoke heat hyperalgesia, and sensitising cold responses in mice (Lippoldt et al. 2013). In DRG neurons, artemin-GFRα3 signaling has been shown to upregulate TRPV1 mRNA via the p38 MAPK pathway (Elitt et al. 2006; Ikeda-Miyagawa et al. 2015) and artemin also upregulates TRPM8 expression (Yang et al. 2024). Artemin has also been shown to upregulate the p-JNK pathway (Zhu et al. 2020), inhibition of which in primary sensory neurons can suppress mechanical allodynia and alleviate pain in rats (Zhuang et al. 2006). Therefore, the p-JNK pathway may underlie the pain progression associated with artemin signaling. The sensitization of TRPV1 by artemin in mouse and NMR DRG neurons observed in this study aligns with what others have observed previously in mouse (Elitt et al. 2006; Malin et al. 2006), and although the signalling mechanisms producing this sensitization are unknown, it would appear that there are conserved mechanisms across mice and NMRs, i.e. in the same way that activation of PKC signalling has been shown to sensitize TRPV1 from a variety of species, including mice and NMRs (Vellani et al. 2001; Correll et al. 2004; Omerbašić et al. 2016). Behaviorally, numerous studies have shown that activation of artemin-GFRα3 signaling correlates with increased pain sensitivity. For example, exogenous administration of artemin to mice leads to heightened pain behaviors and sensitization of nociceptive neurons innervating joints, indicating a direct role in pain perception (Minnema et al. 2022; Morgan et al. 2023). Conversely, blocking artemin signaling in rat model of osteoarthritis with neutralizing antibodies prevents pain and reverses nociceptive neuron hypersensitivity, highlighting its potential as a therapeutic target (Morgan et al. 2023). However, translating these preclinical findings to clinical applications has proven challenging, a recent study failing to show any efficacy of an anti-GFRα3 antibody in patients with osteoarthritic knee pain (Somersan-Karakaya et al. 2023).

Although modulation of TRPV1 function by artemin has previously been investigated in rodent sensory neurons (Elitt et al. 2006; Malin et al. 2011), to our knowledge, this study is the first to investigate the ability of artemin to modulate intrinsic neuronal excitability. We observed that acute exposure of both mouse and NMR DRG neurons to artemin leads to depolarization of the resting membrane potential and a decrease in rheobase, indicating increased neuronal excitability. These findings align with previous reports demonstrating that artemin increases mechanical sensitivity of joint afferents and sensitizes bone afferents (Morgan et al. 2023). Although we cannot rule out any direct effect of artemin-GFRα3 signalling on the function of mechanosensitive ion channels, such as Piezo2 and Elkin1 (Coste et al. 2010; Chakrabarti et al. 2024), the artemin-induced changes in intrinsic excitability likely contribute to effects that others have observed regarding changes in mechanosensitivity. From these data, it would also appear that without any stimulation, NMR DRG neurons have a more depolarized RMP than mouse DRG neurons, further studies being required to confirm this pattern and identify the underlying mechanism. Overall, our results provide further insight into the cellular mechanisms underlying artemin-induced sensitization, suggesting that modulation of intrinsic excitability contributes to the observed behavioral hypersensitivity. Future work will be required to work out the signalling mechanisms subserving the sensitization observed.

In summary, employing an *in vitro* methodology, we observed that artemin sensitizes both the intrinsic electrical properties of mouse and NMR DRG sensory neurons as well as sensitivity to capsaicin, results suggesting the conserved nature of artemin-GFRα3 signaling in sensory neurons across mice and NMRs. Due to the fact that we observed similar artemin-induced sensitization of sensory neurons *in vitro*, we decided not to conduct *in vivo* experiments from a 3Rs perspective, i.e. artemin is likely to induce nociception, and thus suffering and distress, in both species. Consequently, a limitation of our approach is that we cannot be certain that the ability of artemin to sensitize both mouse NMR sensory will result in the ability of artemin to induce hyperalgesia in NMRs as it does in other rodents (Minnema et al. 2022; Morgan et al. 2023).

## Statements and Declarations

The authors declare that they have no competing financial interests or personal relationships that could have influenced the work reported in this paper.

## Acknowledgements

The authors thank the hard work and care provided to naked mole-rats and mice by staff within the Combined Animal Facility at the University of Cambridge. The authors acknowledge support from the Dunhill Medical Trust (RPGF2002\188) and the David James Trust Fund.

## Data availability

Data to support this article are available https://doi.org/10.17863/CAM.117370.

